## Supplementary Figures 1-7 for "HIV-1 Vpr antagonizes innate immune activation by targeting karyopherin-mediated NF-κB/IRF3 nuclear transport"

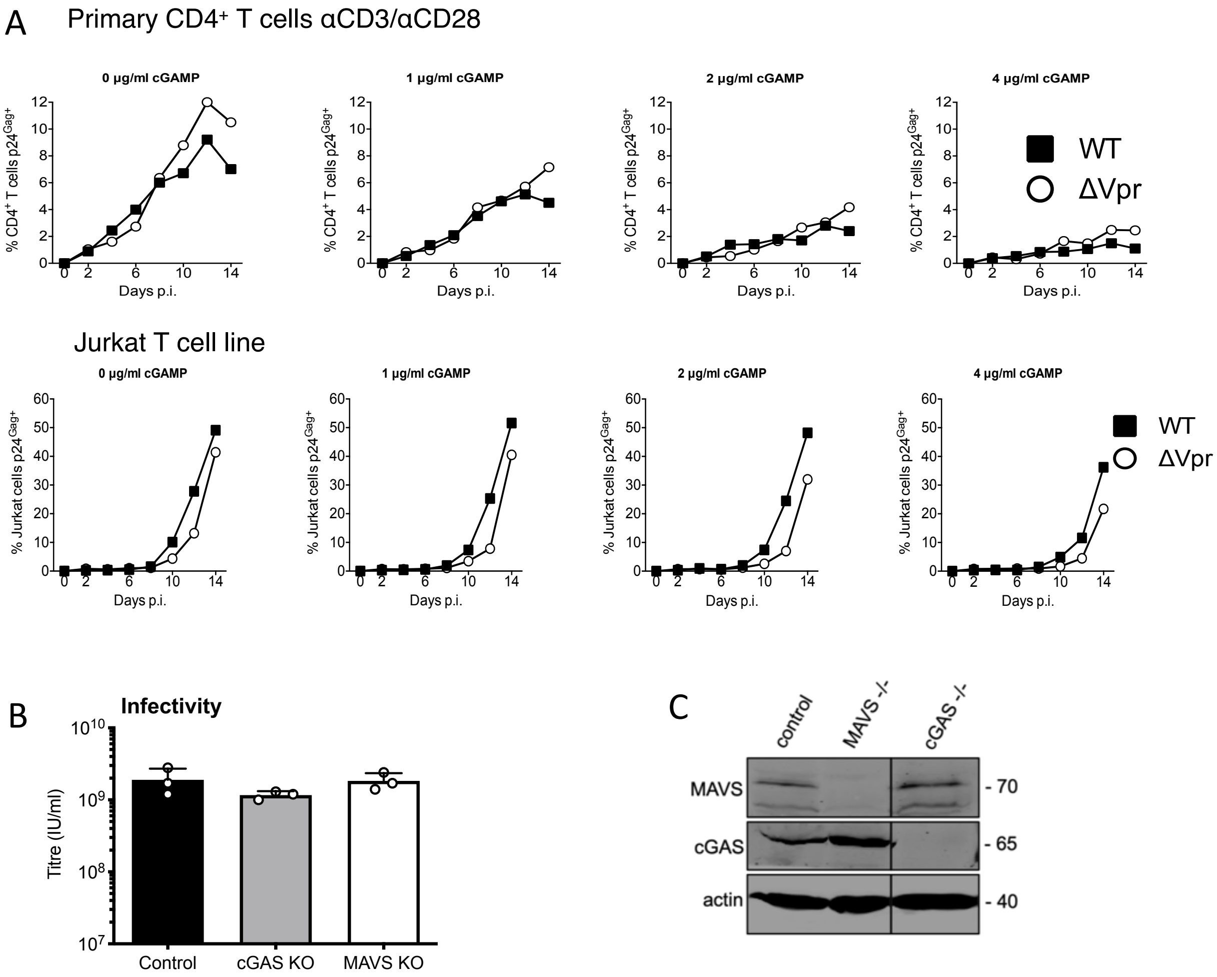


**Figure S1 HIV-1 replication in cGAMP stimulated MDMs requires Vpr and Vpr suppresses HIV-1 innate immune sensing by cGAS**

(**A**) Replication of WT NL4-3 HIV-1 or NL4-3 HIV-1ΔVpr in activated CD4+ T cells stimulated with 1 μg/ml, 2 μg/ml or 4 μg/ml cGAMP or left unstimulated, measured by flow cytometry staining infected cells with anti-p24 antibody. (**B**) HIV-GFP titre in control, cGAS-/- or MAVS-/- THP-1 cells used in Figure 1 (G). (**C**) Immunoblot detecting cGAS, MAVS, or actin as a loading control, from extracted cGAS-/- or MAVS-/- knock out THP-1 cells or their CRISPR/Cas control cells. Size marker positions are shown on the right (kDa).

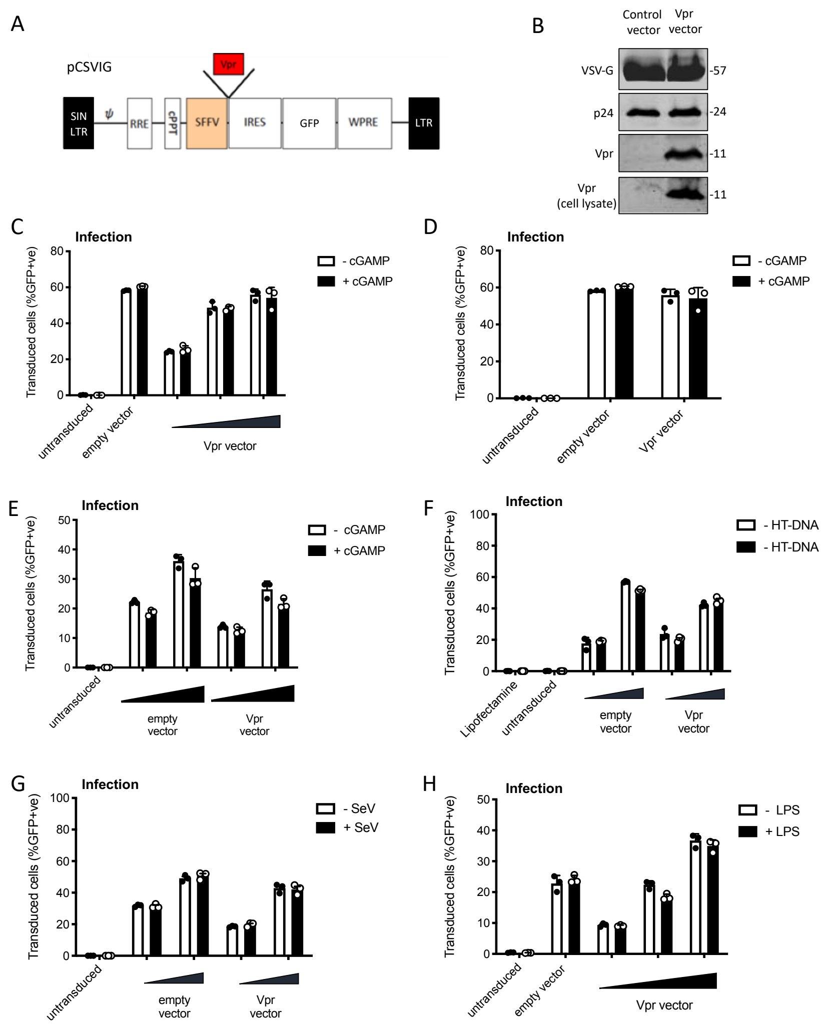


**Figure S2 HIV-1 Vpr expression inhibits interferon stimulated gene expression after stimulation with various innate immune stimuli**

(**A**) Vpr encoding lentiviral expression construct (pCSVIG) contained self-inactivating Long terminal repeat (SIN LTR), Rev response element (RRE), Central polypurine tract (cPPT), Spleen focus-forming virus promoter (SFFV), internal ribosome entry site (IRES), green fluorescent protein (GFP) and Woodchuck hepatitis virus post-transcriptional regulatory element (WPRE). (**B**) Immunoblot detecting VSV-G envelope, capsid (p24) and Vpr in vector supernatant and Vpr additionally in target cell lysate. Size markers in kDa are indicated on the right. (**C**) Percentage of THP-1 cells in Figure 2A transduced by the vector encoding Vpr and GFP (MOI 0.25, 0.5, 1) or empty vector encoding GFP alone (MOI 1) and treated with cGAMP (5 µg/ml) or left untreated as a control. (**D**) Percentage of THP-1 cells in Figure 2B transduced by the vector encoding Vpr and GFP (MOI 1) or empty vector encoding GFP alone (MOI 1) and treated with cGAMP (5 µg/ml) or left untreated as a control. (**E**) Percentage of THP-1 cells in Figure 2C transduced by the vector encoding Vpr and GFP (MOI 0.5, 1) or empty vector expressing GFP alone (MOI 0.5, 1) and treated with cGAMP (5 µg/ml) or left untreated as a control. (**F**) Percentage of THP-1 cells in Figure 2D transduced by the vector encoding Vpr and GFP (MOI 0.5, 1) or empty vector encoding GFP alone (MOI 0.5, 1) and stimulated with HT-DNA transfection (5 µg/ml) or left untreated as a control. (**G**) Percentage of THP-1 cells in Figure 2E transduced by the vector encoding Vpr and GFP (MOI 0.5, 1) or empty vector expressing GFP alone (MOI 0.5, 1) and stimulated with Sendai virus infection or left untreated as a control. (**H**) Percentage of THP-1 cells in Figure 2F transduced by the vector encoding Vpr and GFP (MOI 0.25, 0.5, 1) or empty vector encoding GFP alone (MOI 1) and stimulated with LPS treatment (1 µg/ml) or left untreated as a control.

Data are expressed as means ± SD (n = 3). Data are representative of three (C, F-H) or two (B, D, E) independent experiments.

expressing WT, or mutant, Vpr from a lentiviral vector (MOI 1), or after transduction by empty vector (MOI 1) or in untransduced THP-1. Data are mean ± SD (n = 3). Two-way ANOVA: * ($p<0.05$), ** ($p<0.01$), *** ($p<0.001$), **** ($p<0.0001$) compared to no Vpr or empty vector controls. Data are representative of three (B-D, F) or two (A, E, G) independent experiments.

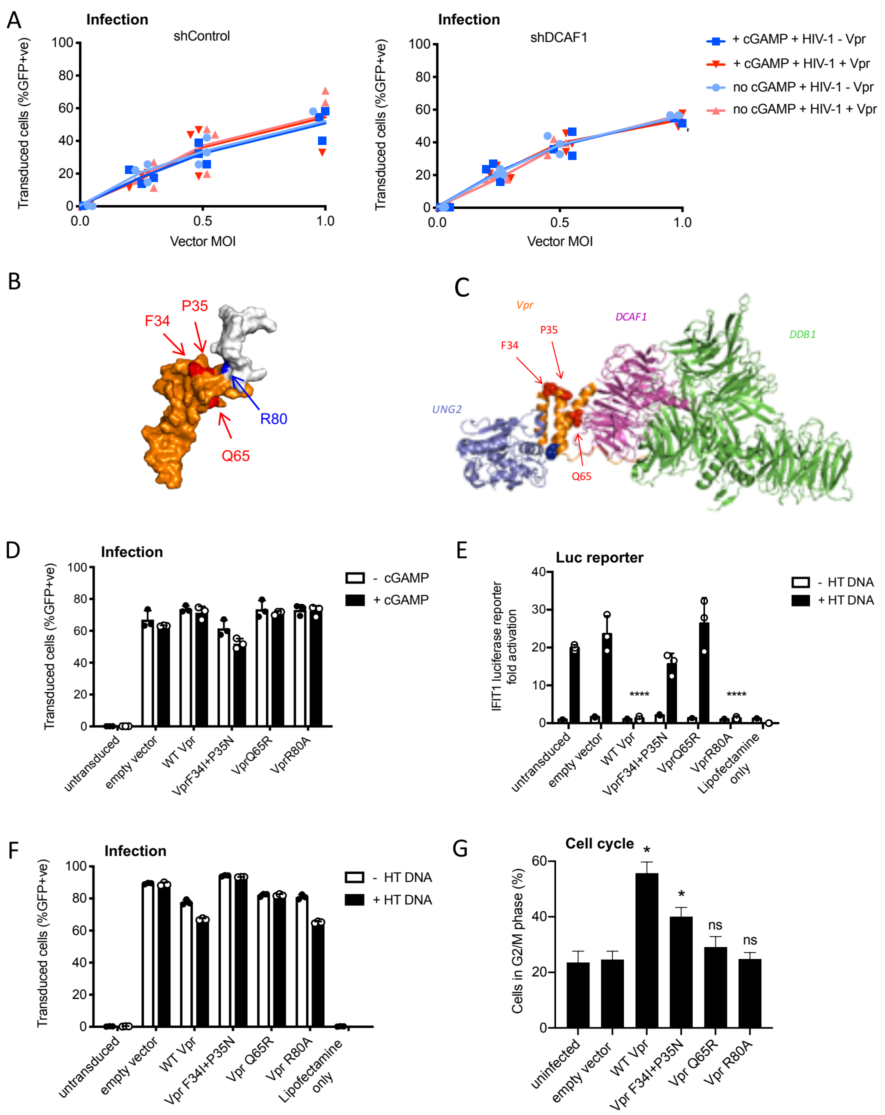


**Figure S3 Vpr inhibition of innate immune activation is dependent on DCAF1 but independent of cell cycle arrest**

(**A**) Percentage of THP-1 cells in Figure 3C transduced by the vector encoding Vpr and GFP, or empty vector encoding GFP alone, at the indicated MOI and treated with cGAMP (5 µg/ml) or left untreated. (**B**) NMR structure of full length Vpr showing position of Vpr mutants (PDB: 1M8L). White region (c-terminus) of Vpr shown in (B) is unresolved in the crystal structure (C). (**C**) Crystal structure of Vpr (orange) with its target protein UNG2 (blue) and cofactors DCAF1(pink) and DDB1 (green) showing position of Vpr mutations (PDB: 5JK7). (**D**) Percentage of THP-1 cells in Figure 3F transduced by the vector encoding WT, or mutant, Vpr and GFP (MOI 1), or empty vector encoding GFP alone (MOI 1), and treated with cGAMP (5 µg/ml), or left untreated as a control. (**E**) Fold induction of IFIT1-Luc after HT-DNA (5 µg/ml) transfection in cells expressing WT, or mutant, Vpr from a lentiviral vector (MOI 1), or empty vector (MOI 1), or in untransduced IFIT1-Luc reporter THP-1 cells. (**F**) Percentage of THP-1 cells in Figure S3E transduced with HIV-1 vector encoding WT, or mutant, Vpr and GFP (MOI 1), or empty vector encoding GFP alone (MOI 1), and transfected with HT-DNA (5 µg/ml) or left untransfected as a control. (**G**) Percentage of THP-1 cells in G2/M phase of cell cycle after transduction with an empty vector (MOI), or vector encoding WT Vpr, or mutant Vpr, (MOI 1) or left untransduced as a control. Mean+/- SEM n=2.
Unless stated data are expressed as means ± SD (n = 3). Data is analysed using two-way ANOVA test. * ($p<0.05$), ** ($p<0.01$), *** ($p<0.001$), **** ($p<0.0001$) compared to empty vector. Data are representative of three (A), (D) or two (E-G) independent experiments.

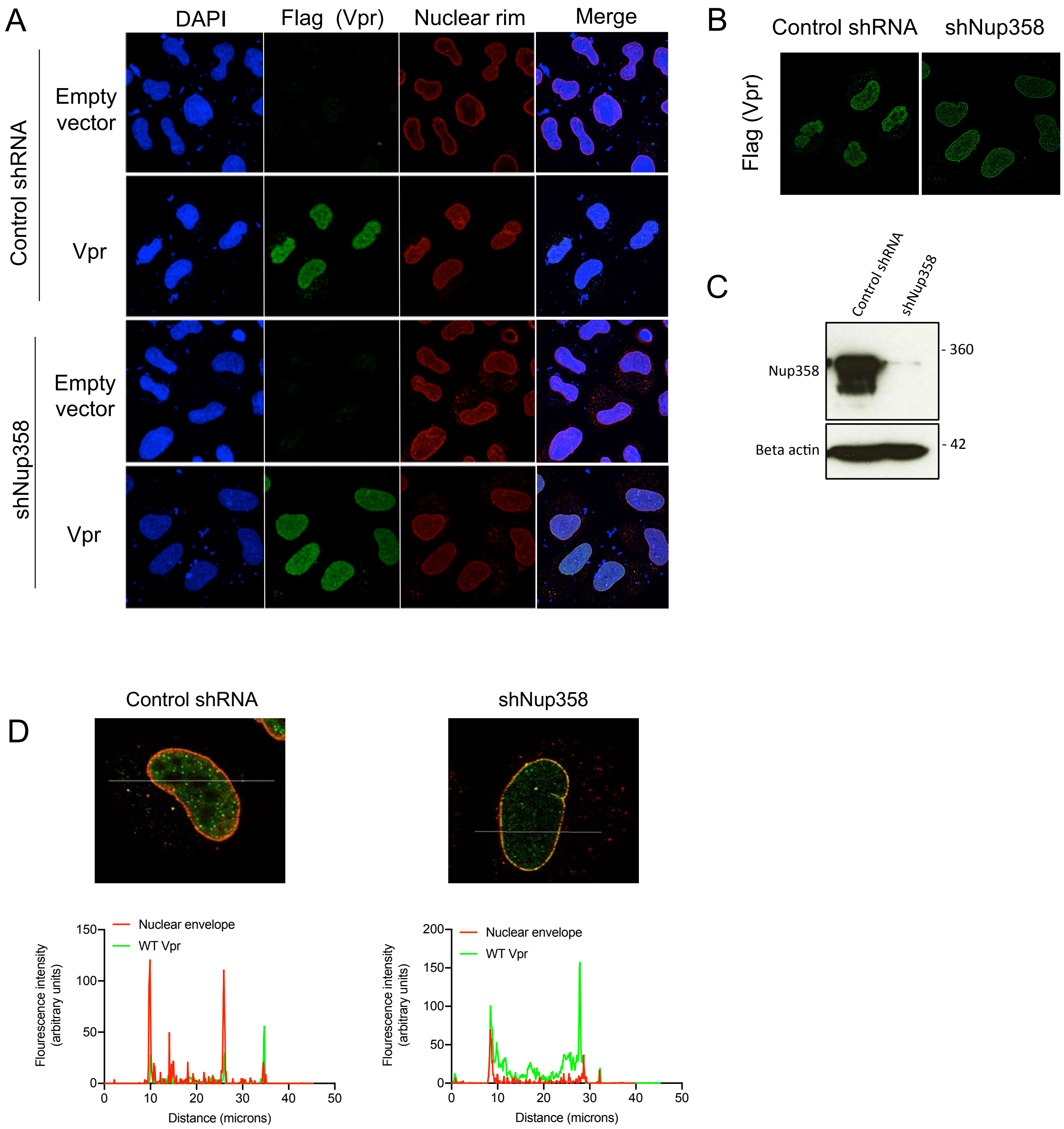


**Figure S4 Nup358 is not required for Vpr colocalization with mab414 nuclear pore staining**

(**A**) Immunofluorescence images of HeLa cells expressing a control, or Nup358 targeting, shRNA transfected with empty vector or Flag-tagged Vpr encoding pcDNA3.1 plasmid (50 ng) using antibodies detecting the Flag-tag (green) or the nuclear pore complex (mab414) (red). 4′,6-Diamidine-2′-phenylindole dihydrochloride (DAPI) stains nuclear DNA (Blue). (**B**) Selected confocal images (z-section) of cells in (A) showing effect of Nup358 depletion on colocalization of Vpr with mab414 nuclear pore staining (**C**) Immunoblot detecting Nup358, or actin as a loading control, from extracted Hela cells expressing a control, or Nup358 targeting, shRNA in cells from A. Size markers are shown (kDa). (**D**) Assessment of colocalization of Flag-tagged Vpr and mab414 stained nuclear pores in cells expressing a control, or Nup358 targeting, shRNA.

immunofluorescence measurement of IRF3 nuclear translocation in PMA differentiated THP-1 cells stimulated with cGAMP (5 µg/ml), or left unstimulated, and then immediately infected with HIV-1 GFP lacking Vpr or bearing WT Vpr or Vpr mutants as shown (1 RT U/ml) or left uninfected. (**G**) Single cell immunofluorescence measurement of IRF3 nuclear translocation in PMA differentiated THP-1 cells transfected with HT-DNA (5 µg/ml), or left untransfected, and immediately infected with HIV-1 GFP lacking Vpr, or bearing WT or mutant Vpr (1 RT U/ml) or left uninfected.

Data in B is expressed as means ± SEM (n = 2). Data is analysed using two-way ANOVA: * ($p<0.05$), ** ($p<0.01$), *** ($p<0.001$), **** ($p<0.0001$) compared to data from infection with HIV-1 lacking Vpr. Data are representative of three (C–G) or two (A, B) independent experiments.

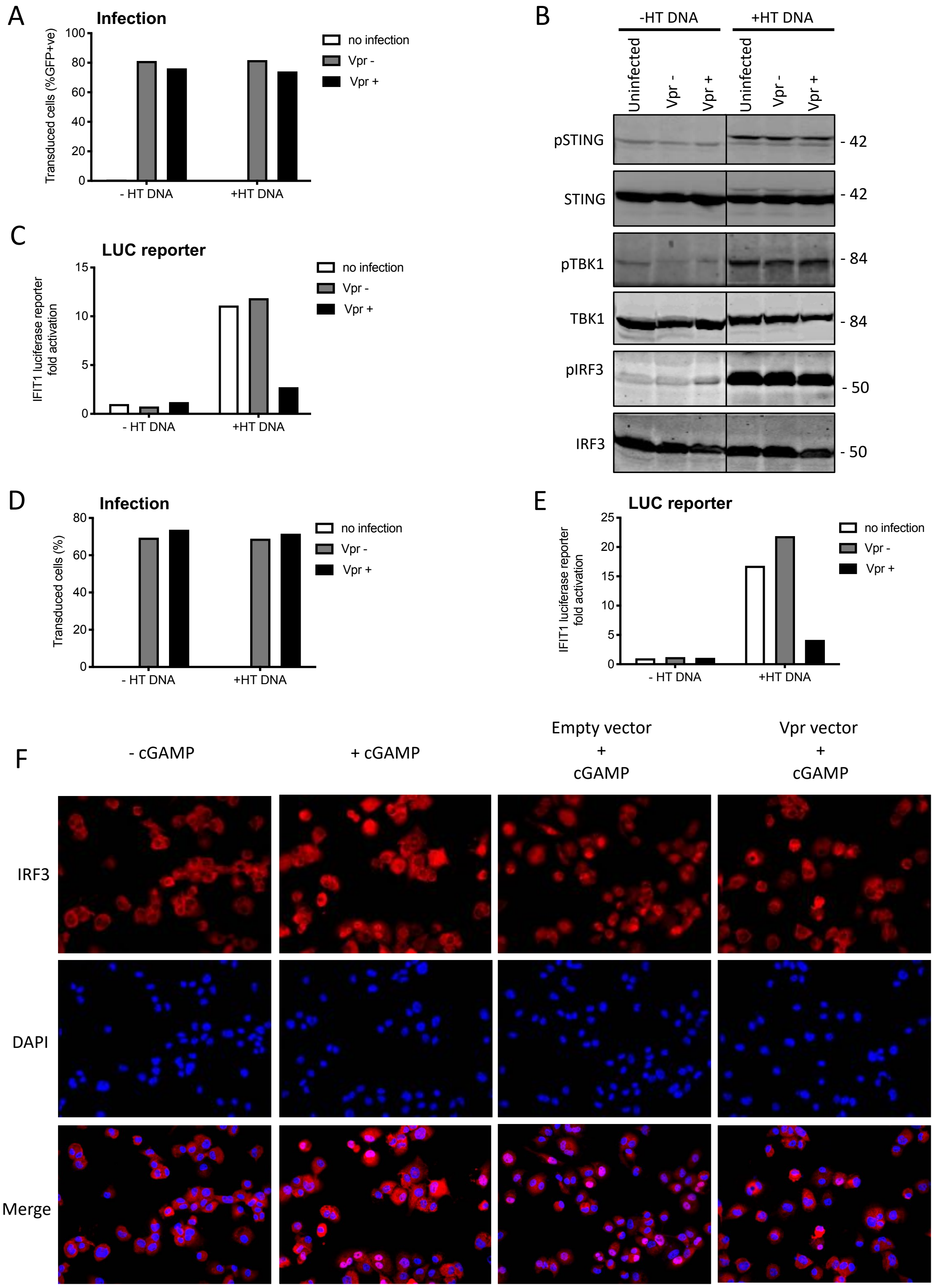
A
Infection
Transduced cells (%GFP+ve)
100
80
60
40
20
0
- HT DNA
+HT DNA
no infection
Vpr -
Vpr +
B
-HT DNA
+HT DNA
Uninfected
Vpr -
Vpr +
Uninfected
Vpr -
Vpr +
pSTING
- 42
STING
- 42
pTBK1
- 84
TBK1
- 84
pIRF3
- 50
IRF3
- 50
C
LUC reporter
IFIT1 luciferase reporter fold activation
15
10
5
0
- HT DNA
+HT DNA
no infection
Vpr -
Vpr +
D
Infection
Transduced cells (%)
80
60
40
20
0
- HT DNA
+HT DNA
no infection
Vpr -
Vpr +
E
LUC reporter
IFIT1 luciferase reporter fold activation
25
20
15
10
5
0
- HT DNA
+HT DNA
no infection
Vpr -
Vpr +
F
- cGAMP
+ cGAMP
Empty vector + cGAMP
Vpr vector + cGAMP
IRF3
DAPI
Merge

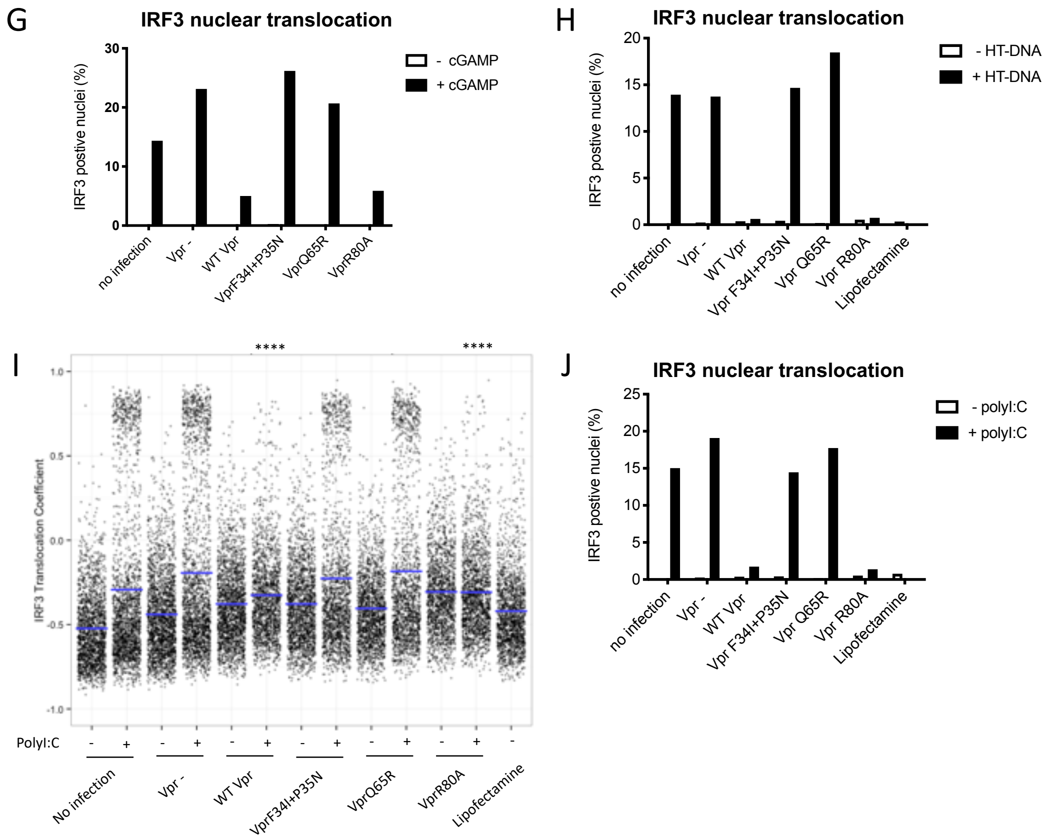


**Figure S5 Vpr inhibits IRF3 nuclear translocation**

(**A**) Percentage of THP-1 cells in Figure 5A transduced by HIV-1 GFP vector bearing Vpr , or HIV-1 GFP lacking Vpr, transfected with HT-DNA (5 µg/ml) or left untransfected as a control. (**B**) Immunoblot detecting Phosho-STING (Ser366), total STING, phospho-TBK1 (Ser172), total TBK1, phospho-IRF3 (Ser386) or total IRF3 from extracted THP-1 cells expressing Vpr, empty vector or left untransduced as a control, and transfected with HT-DNA (5 µg/ml), or left untransfected as a control. Size markers are shown (kDa). (**C**) Fold induction of IFIT1-Luc in cells from gel in Figure 5A, expressing Vpr, or empty vector, and transfected with HT-DNA (5 µg/ml) or left untransfected as a control. (**D**) Percentage of THP-1 cells from Figure S5B transduced by HIV-1 GFP bearing Vpr, or lacking Vpr, transfected with HT-DNA (5 µg/ml) or left untransfected as a control. (**E**) Fold induction of IFIT1-Luc in cells from second experiment (gel presented in Figure S5B) expressing Vpr, or empty vector, and transfected with HT-DNA (5 µg/ml) or left untransfected as a control. (**F**) Representative immunofluorescence images showing IRF3 (red) nuclear translocation in PMA differentiated THP-1 cells treated with cGAMP, or left untreated, and infected with HIV-1 GFP bearing Vpr, or lacking Vpr, or left uninfected. 4′,6-Diamidine-2′-phenylindole dihydrochloride (DAPI) stains nuclear DNA (Blue). (**G**) Percentage of cells with IRF3 translocation coefficient greater than 0.5 plotted as a percentage from Figure 5F. (**H**) Percentage of cells with IRF3 translocation coefficient greater than 0.5 plotted as a percentage from Figure 5G. (**I**) Single cell measurement of IRF3 nuclear translocation in PMA differentiated THP-1 cells transfected with poly I:C, or left untransfected, and infected with HIV-1 GFP lacking Vpr or bearing WT or mutant Vpr (1 RT U/ml), or left uninfected. (**J**) Percentage of cells with IRF3 translocation coefficient greater than 0.5 plotted as a percentage from Figure S5I. Data is analysed using two-way ANOVA test: * ($p<0.05$), ** ($p<0.01$), *** ($p<0.001$), **** ($p<0.0001$) compared to empty vector. Data are representative of three (F-J) or two (A-E) independent experiments.

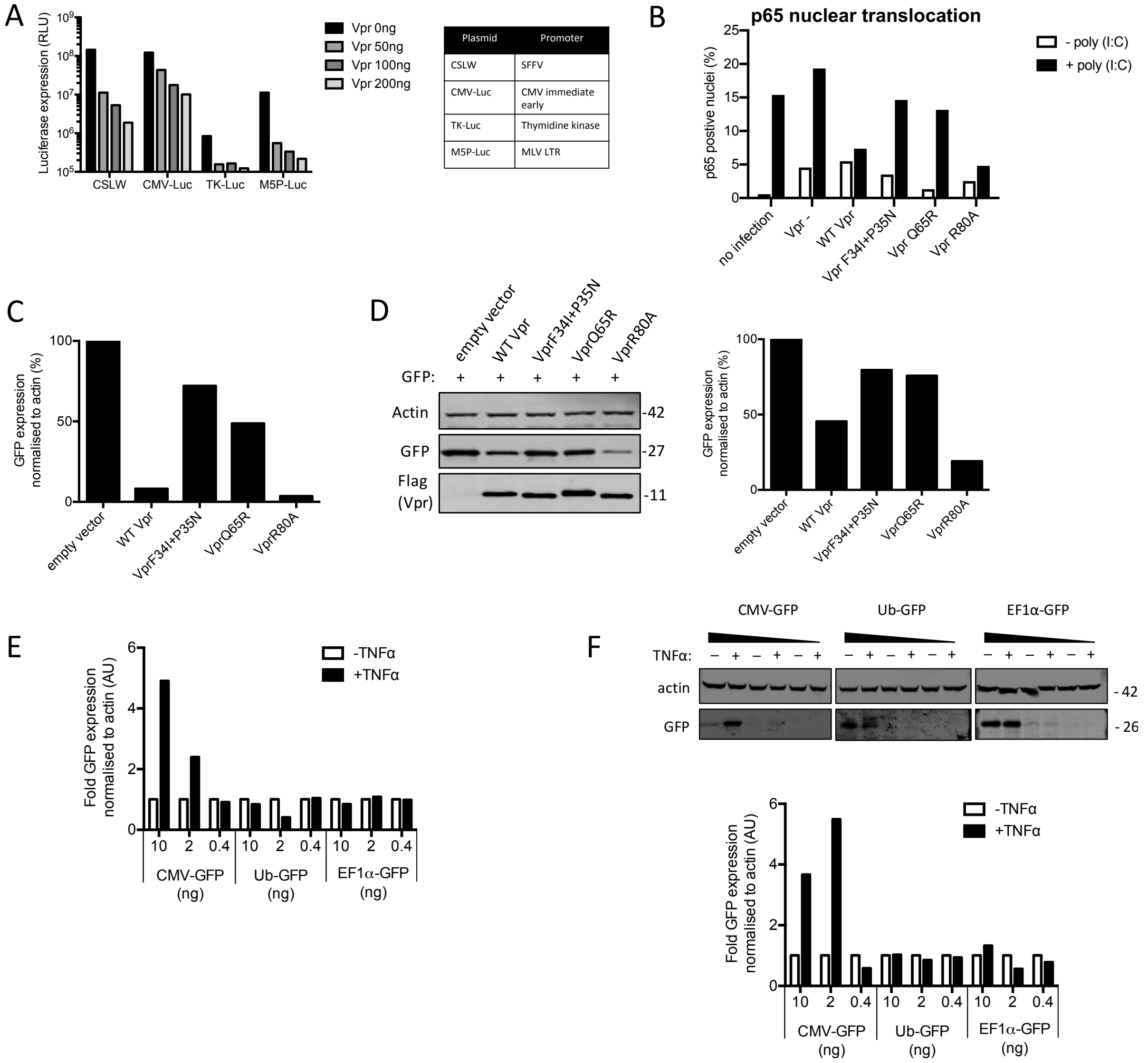


**Figure S6 Vpr inhibits NF-κB p65 nuclear translocation and NF-κB sensitive plasmid expression**

(**A**) Induction of luciferase reporter in HEK293T cells transfected with CSLW, CMV-Luc, TK-Luc or M5P-Luc (10ng), and empty vector, or Vpr encoding vector (50 ng, 100 ng, 200 ng). Table shows the promoters driving the luciferase reporter in each plasmid. (**B**) Percentage of cells in Figure 6D with translocation coefficient greater than 0.5. (**C**) Quantification of GFP expression by densitometry for the immunoblot in Figure 6E. (**D**) Immunoblot detecting flag-Vpr, GFP or actin as a loading control from HEK293T cells transfected with empty vector, flag-tagged WT Vpr encoding vector or flag-tagged mutant Vpr encoding vector and CMV-GFP vector or left untransfected. Size markers are shown in kDa. Quantification of GFP expression by densitometry for the immunoblot is shown on the right. (**E**) Quantification of GFP expression by densitometry for the immunoblot in Figure 6G. (**F**) Immunoblot detecting GFP, or actin as a loading control, from HEK293T cells transfected with CMV-GFP, EF1α-GFP or Ub-GFP plasmids (10 ng, 2 ng, 0.4 ng) and stimulated with TNFα (200 ng/ml) or left unstimulated. Size markers are shown in kDa. Quantification of GFP expression by densitometry for the immunoblot is shown in the right.

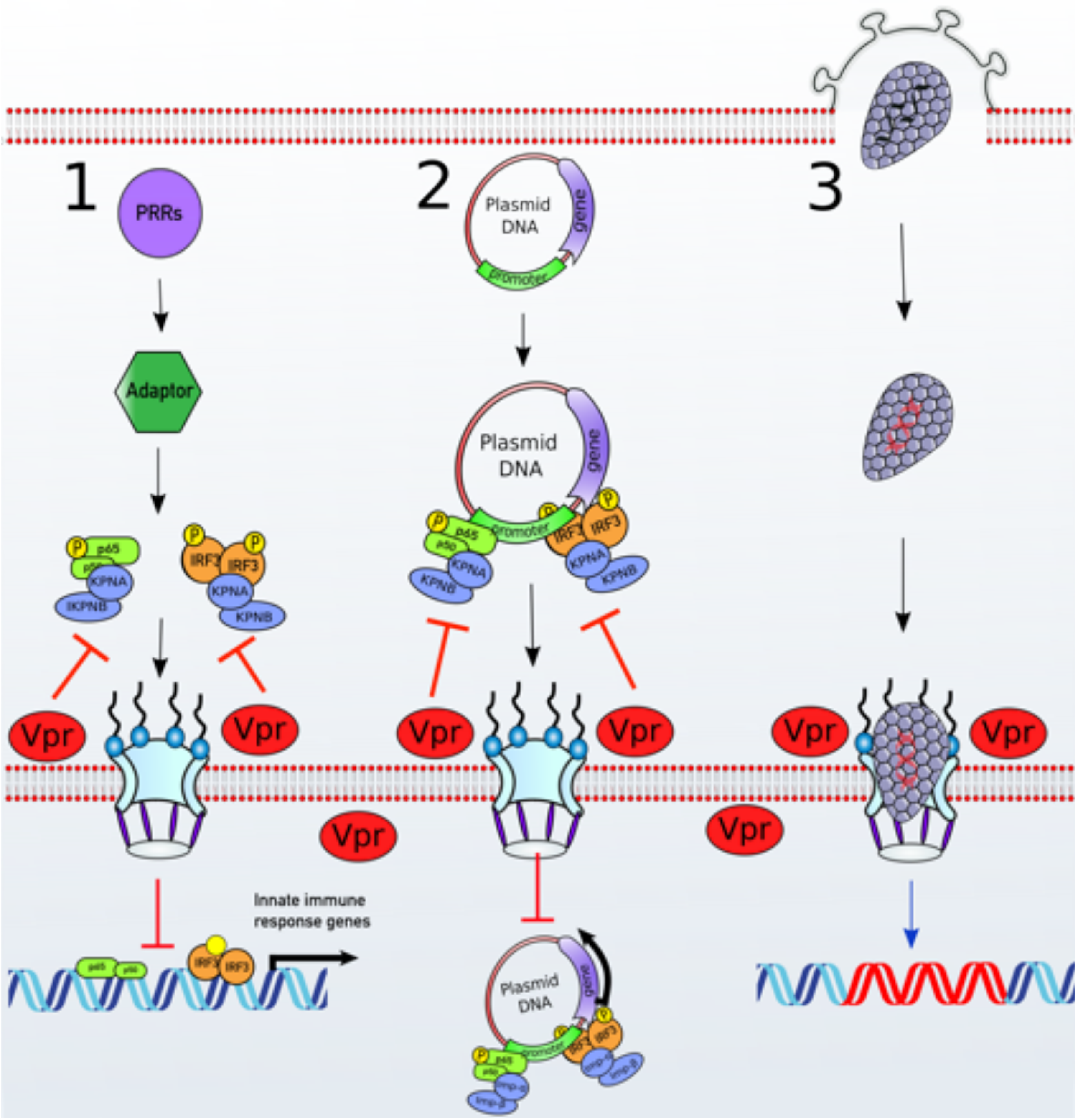


**Figure S7 A unifying model of Vpr function**

**(1)** Stimulation of various PRRs results in activation of transcription factors such as IRF3 and NF-κB. To activate ISGs or proinflammatory genes expression, NF-κB and IRF3 translocate to the nucleus via the classical Karyopherin-α/β dependent nuclear import pathway. **(2)** Nuclear import of a plasmid transfected into cellular cytoplasm is essential for gene expression. Transcription factors such as IRF3 and NF-κB bind to their cognate response elements present in the promoter of the plasmid and allow nuclear import via the classical karyopherin-α/β dependent pathway (Mesika et al., 2001) as well as transcription. **(3)** HIV-1 based vectors deliver genes to the nucleus in a karyopherin-α/β independent manner. Vpr localises to the nuclear pores and targets karyopherin-α dependent nuclear import in a DCAF1 E3 ubiquitin ligase dependent manner. This inhibits nuclear translocation of transcription factors such as IRF3 and NF-κB and subsequent antiviral ISG expression. This also inhibits IRF3 and NF-κB dependent plasmid expression or nuclear import but does not impact lentiviral gene delivery.
